# UMINT: Unsupervised Neural Network For Single Cell Multi-Omics Integration

**DOI:** 10.1101/2022.04.21.489041

**Authors:** Chayan Maitra, Dibyendu Bikash Seal, Vivek Das, Rajat K. De

**Affiliations:** Machine Intelligence Unit, Indian Statistical Institute, 203 Barrackpore Trunk Road, Kolkata 700108, India; A. K. Choudhury School of Information Technology, University of Calcutta, JD-2, Sector - III, Saltlake, Kolkata - 700106, India; Novo Nordisk A/S, Novo Nordisk Park 1, 2760 Maløv, Denmark

**Keywords:** Single cell analyses, Deep neural network, Unsupervised learning, CITE-seq, Multi-omics integration

## Abstract

Multi-omics studies have enabled us to understand the mechanistic drivers behind complex disease states and progressions, thereby providing novel and actionable biological insights into health status. However, integrating data from multiple modalities is challenging due to the high dimensionality of data and noise associated with each platform. Non-overlapping features and technical batch effects in the data make the task of learning more complicated. Conventional machine learning (ML) tools are not quite effective against such data integration hazards. In addition, existing methods for single cell multi-omics integration are computationally expensive. This has encouraged the development of a novel architecture that produces a robust model for integration of high-dimensional multi-omics data, which would be capable of learning meaningful features for further downstream analysis. In this work, we have introduced a novel Unsupervised neural network for single cell Multi-omics INTegration (UMINT). UMINT serves as a promising model for integrating variable number of single cell omics layers with high dimensions, and provides substantial reduction in the number of parameters. It is capable of learning a latent low-dimensional embedding that can capture useful data characteristics. The effectiveness of UMINT has been evaluated on benchmark CITE-seq (paired RNA and surface proteins) datasets. It has outperformed existing state-of-the-art methods for multi-omics integration.

## 1 Introduction

Recent advancements in single cell technologies have provided unprecedented opportunities in analysis of omics data. This allows researchers to probe biological functions at the cellular level while studying embryonic development, immune system or cancer [1, 2, 3]. Existing technologies include DROP-seq [4], SMART-seq2 [5] and 10x Genomics, which allow measuring mRNA expressions at single cell resolution (scRNA-seq). Most recently, technologies have further scaled up to produce data assays from multiple modalities. This has provided several views of the same cell of interest, thereby refining our definitions of the cellular identity. A few such methods include CITE-seq [6] and REAP-seq [7] which enable paired measurement of RNA and cell surface proteins. Other methods, such as ATAC-seq [8], SNARE-seq [9], sci-CAR [10] and SHARE-seq [11] measure paired gene expression and chromatin accessibility. scNMT [12], on the other hand, integrates single cell chromatin accessibility, DNA methylation and transcriptomics data. However, integration of various modalities of data does not come without its challenges [13]. High data dimension, sensitivity associated with each platform, zero-inflation due to dropouts [14], non-overlapping features and batch effects account for the stochasticity and noise in the data.

At present, several methods exist that can perform the task of integration of single cell omics modalities. Seuratv3 [15] can integrate various single cell omics datasets including RNA-seq, protein expression, chromatin and spatial data, and transfer information between them. Seuratv4 [16] provides multi-modal single cell analysis using “weighted nearest neighbor” method. MOFA+ [17], based on Bayesian Group Factor Analysis, is another method that generates a low-dimensional representation of the data by integrating two or more omics among gene expression, DNA methylation and chromatin data. Neural network based methods, like TotalVI [18], uses an encoder function to learn a joint representation of the data and Bayesian inference to build a latent embedding from single cell RNA and protein expressions. Multigrate [19] develops an alternative pipeline for integrating CITE-seq and single cell ATAC-RNA data for both paired and unpaired samples. It has been used to map multimodal queries to reference atlases and impute missing values. Other standard omics integration methods include UINMF [20], MUON [21], scMOC [22] and SIMBA [23]. A comprehensive review of major single cell multi-omics integration methods can be found at [24].

Although different omics layer measurements are recorded against the same set of cells, they might encode different underlying transcriptional states and activities. Existing methods to handle these datasets are not always capable of extracting features relevant to a biologically significant problem. An inherent feature of multi-omics data in concern is its complex, non-linear, layered structure. The architecture of a deep neural network also resembles such layered non-linearity. The output from each layer is multiplied by its weight vector to compute the weighted sum, and a non-linear function is then applied over the weighted sum for each node in the layer. The non-linear output is then passed on to the next layer. Thus, deep learning models facilitate learning complex features in an unsupervised manner. However, existing neural network based methods for single cell multiomics integration are computationally expensive since they involve substantial amount of parameter training. Other methods make assumptions about data distribution, which is unrealistic.

Thus, driven by the need to develop a robust integration method for single cell multi-omics integration, in this work, we have introduced a novel Unsupervised neural network for single cell Multi-omics INTegration (UMINT). UMINT is capable of extracting relevant features from different single cell omics layers. It does not make assumptions about the distribution of data and is competent enough to integrate variable number of omics modalities. In addition, UMINT is computationally far less expensive than other existing neural network based methods used for single cell multi-omics integration. The performance of UMINT has been evaluated on four publicly available CITE-seq datasets. Results compare favourably against other existing state-of-the-art methods, which makes UMINT completely fit in with the status quo.

## 2 Methodology

This section describes the datasets used in the experiments, the methodology used for data pre-processing and the proposed neural network model, called UMINT, for single cell multi-omics integration.

### 2.1 Data acquisition and pre-processing

Four publicly available CITE-seq datasets, viz., *cbmc*8*k* [25], *pbmc*10*kMALT* [26], *bmcite*30*k* [15] and *kotliarov*50*k* [27] have been used in this work. The datasets have been downloaded as count matrices and preprocessed via Seuratv4 [16]. For scRNA-seq part in these datasets, we have normalized them by library size to sum up to 10000, applied a logarithmic transformation, extracted highly variable genes, and finally scaled them linearly (with default parameters). The protein expressions/antibody-derived tag (ADT) datasets have been normalized using the centered log-ratio transformation [25]. Three proteins *CCR*5, *CCR*7 and *CD*10 have been removed from *cbmc*8*k* dataset due to poor abundance. The first three datasets, viz., cbmc8k, *pbmc*10*kMALT* and *bmcite*30*k*, have been used to evaluate the performance of UMINT and compare the results against other state-of-the-art algorithms. The fourth dataset, viz., *kotliarov*50*k*, contains filtered cells with highly variable genes only. It has been preprocessed via Seuratv4 and used to assess other performance criteria of the proposed methodology. The summary of the single cell multi-omics datasets used in this work have been listed in Table 1.

**Table 1:**
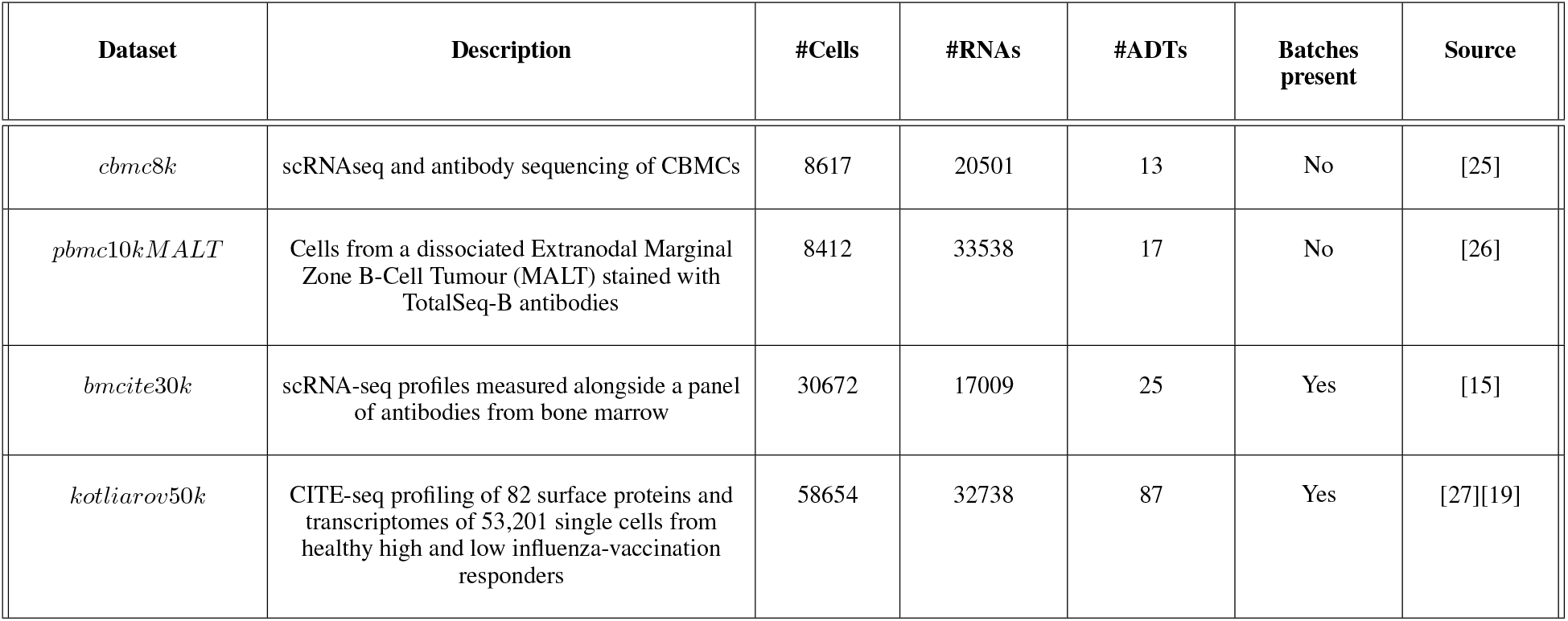
Summary of datasets used for evaluation of the methodology

| Dataset | Description | #Cells | #RNAs | #ADTs | Batches present | Source |
| --- | --- | --- | --- | --- | --- | --- |
| *cbmc8k* | scRNAseq and antibody sequencing of CBMCs | 8617 | 20501 | 13 | No | [25] |
| *pbmc10k MALT* | Cells from a dissociated Extranodal Marginal Zone B-Cell Tumour (MALT) stained with TotalSeq-B antibodies | 8412 | 33538 | 17 | No | [26] |
| *bmcite30k* | scRNA-seq profiles measured alongside a panel of antibodies from bone marrow | 30672 | 17009 | 25 | Yes | [15] |
| *kotliarov50k* | CITE-seq profiling of 82 surface proteins and transcriptomes of 53,201 single cells from healthy high and low influenza-vaccination responders | 58654 | 32738 | 87 | Yes | [27][19] |

### 2.2 Unsupervised neural network for single cell Multi-omics INTegration (UMINT)

In this work, we have developed a deep Unsupervised neural network for single cell Multi-omics INTegration (UMINT). UMINT is a non-recurrent feed-forward neural network that is efficient enough to integrate variable number of omics layers and extract a latent embedding at a reduced dimension. The network structure of UMINT represents a novel neural network architecture as shown in Figure 1.

**Figure 1:**
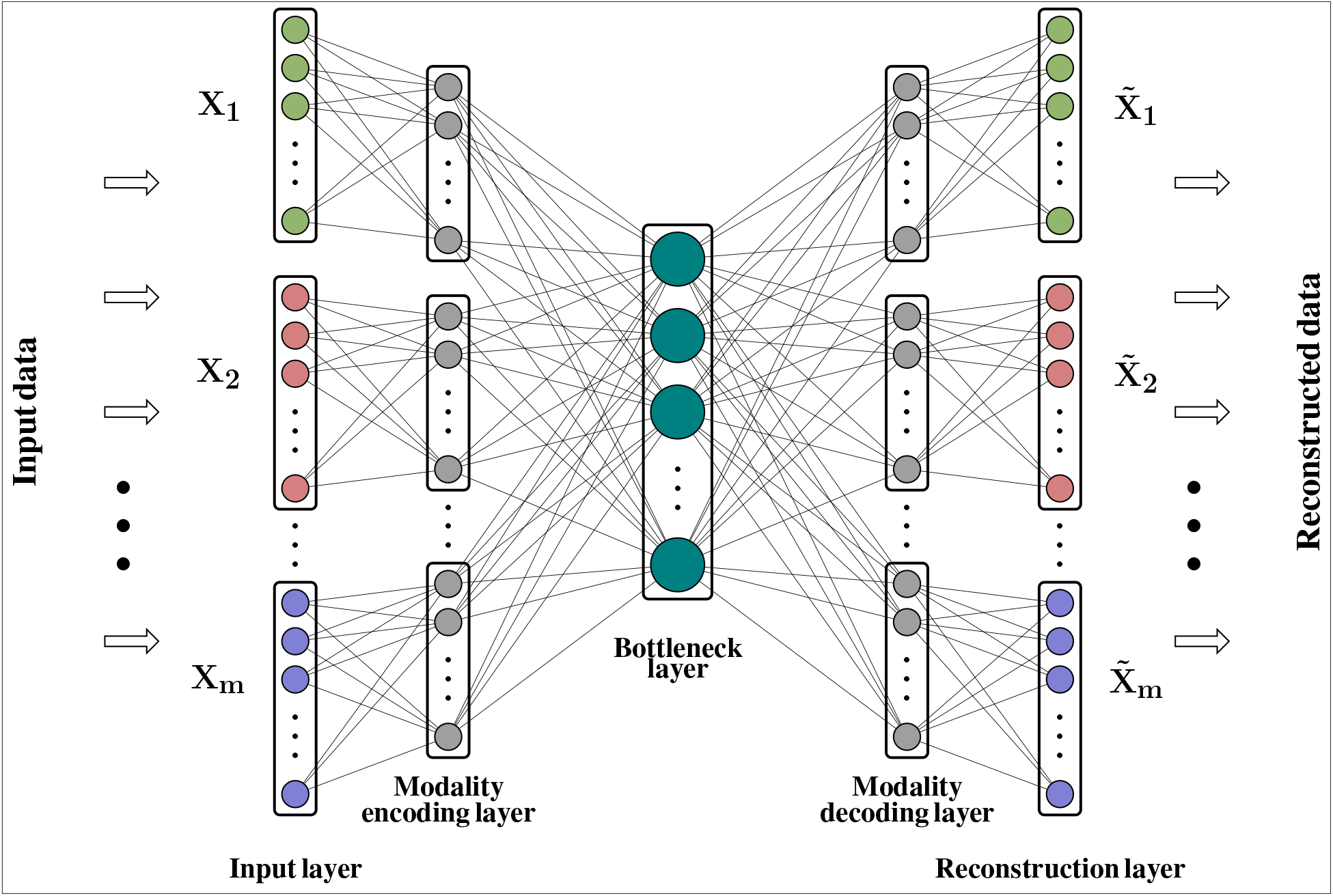
UMINT network structure

Let **X**_1_, **X**_2_,…, **X**_*m*_ be *m* datasets corresponding to *m* different omics modalities having *n* samples each with *d*_1_, *d*_2_, …, *d_m_* features respectively. The UMINT architecture consists of two sub-architectures - an encoder and a decoder. The encoder accepts data from multi-omics datasets presented to the *Input layer,* transports them through one or more *Modality encoding layer*(s) and integrates them in the final layer, known as the *Bottleneck layer.* The decoder accepts the embedded output from the *Bottleneck layer*, transports them through one or more *Modality decoding layer*s and finally tries to reconstruct the original data at the *Reconstruction layer.* In this work, we have used only one layer each for modality encoding and decoding. However, UMINT may contain multiple such layers based on the requirement. In order to improve generalization capability and reduce dimension of the latent embedding, the number of neurons at the *Bottleneck layer* has been kept smaller than the number of neurons in the *Input layer*.

#### 2.2.1 Forward propagation

The *Input layer* of UMINT consists of *m* different modules, each of which accepts input from a data modality. The number of neurons in each of the modules in the *Input layer* is equal to the dimensions of the individual data modalities *d*_1_, *d*_2_,…, *d_m_* respectively. In this layer, UMINT tries to find a suitable projection for each of the data modalities that may be good enough to get integrated in subsequent layers. Each module in the first *Modality encoding layer* shares a dense connection with the corresponding modules of the *Input layer*. The first *Modality encoding layer* containing m modules thus accepts data from m modules in the *Input layer* as input, and obtains m different projections. Let **a**_*ii*_ and **h**_*ji*_ be the input to and the output from the *i^th^* module in the *j^th^* layer respectively. Then, for the *Input* and *Modality encoding* layers, we have

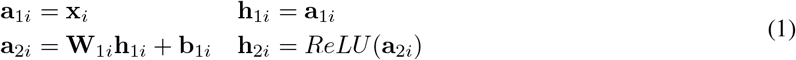

where **x**_i_ is a sample in *i^th^* modality. The term **W**_1*i*_ denotes the weights between *i^th^* module of the *Input layer* and *i^th^* module of the first *Modality encoding layer,* and *b*_1*i*_ denotes the bias terms of the nodes in *i^th^ Modality encoding layer*. Finally, the outputs from the *Modality encoding layer*(s) are projected onto a lower dimensional space in the *Bottleneck layer*. The final *Modality encoding layer* and the *Bottleneck layer* are fully connected. If **W**_2*i*_ represents the weights between the *i^th^* module of the final *Modality encoding layer* and the *Bottleneck layer,* then we have

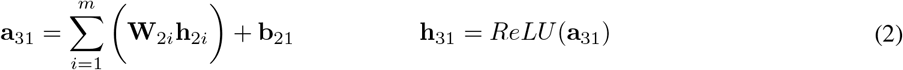

where **b**_21_ denotes the bias terms of the nodes in *Bottleneck layer.* This concludes the process of encoding. The overall function of the encoder network can thus be represented as

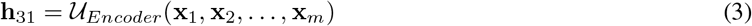

Reconstruction of the original data is done by the decoder network in exactly the opposite manner to that of encoding. The integrated embedding coming out of the *Bottleneck layer* is projected onto the *Modality decoding layer*(s) which consists of the same number of modules as that in the *Modality encoding layer*(s). The number of neurons in each module of the *Modality decoding layer* is identical to that used in the modules in the *Modality encoding layer*. The last layer of the decoder is the *Reconstruction layer* which tries to reconstruct the original data from the respective modules in the final *Modality decoding layer.* The process of decoding can be expressed as

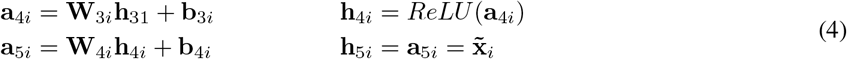

where **W**_*ji*_ and **b**_*ji*_ represent weights and biases for the *i^th^* module in the *j^th^* layer respectively. Thus, the decoder function is given by

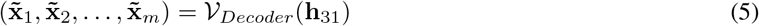

where 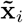 denotes the reconstruction for **x**_*i*_.

#### 2.2.2 Objective function

In this scope of work, UMINT has been used to integrate scRNA-seq and single cell protein expression data. Thus, for each dataset, paired RNA and ADT assays form the inputs to UMINT. For each cell **x**, UMINT tries to find an optimal reconstruction 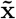 of the input data retaining as much information as possible, thereby minimizing the reconstruction error 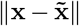. In order to avoid over-fitting, we have used an *L*1 regularization on the nodes’ activities to allow sparsity of nodes’ outputs and an *L*2 regularization on the edge weights since *L*2 regularization tries to shift weight values towards zero. Both *L*1 and *L*2 regularizations minimize the model complexity. In this work, *L*1 and *L*2 regularizations have been controlled using regularization parameters *α* and *β* respectively. The objective function thus becomes

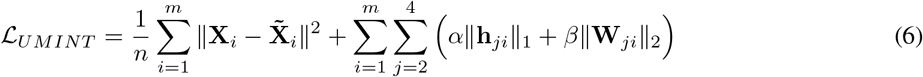

Values of *α* and *β* have been set to 0.0001 and 0.001 respectively, as recommended in literature [28]. UMINT has been trained for 25 epochs using Adam Optimizer [29] and Mean Squared Error (MSE) as the loss function. During the forward pass, the data is fed as input to the encoder. A lossy reconstruction of the input data is produced by the decoder at the *Reconstruction layer.* The error value is then propagated backwards, and the weights and biases are updated for a better reconstruction in the next forward pass.

### 2.3 Latent low-dimensional embedding and clustering

The UMINT model proposed in this work has been used to integrate RNA and protein expression data. Once trained to reconstruct the input data, UMINT is capable of learning a latent low-dimensional embedding that captures meaningful features from the integrated data. This latent embedding has been used in the subsequent step for downstream analysis in order to explore its effectiveness. We have used agglomerative hierarchical clustering and k-means clustering algorithm on this latent embedding to cluster the cell-types for each of the datasets used in the study. The performance of UMINT has then been validated against existing benchmark methods used for multi-omics integration.

## 3 Results

UMINT is an unsupervised neural network model that has the potential to learn significant features from the data. The performance of UMINT has been benchmarked on three datasets *cbmc*8*k*, *pbmc*10*kMALT* and *bmcite*30*k* used in this work, over multiple steps. The latent embedding produced by UMINT has been first compared with the latent embedding produced by another existing unsupervised neural network architecture. Subsequently, the UMINT-generated embedding has been compared with other state-of-the-art single cell multi-omics integration methods. Finally, UMINT has been further tested for its performance on a dataset *kotliarov*50*k* having multiple batches. For evaluating the batch correction performance of UMINT, we have performed two different experiments on *kotliarov*50*k* dataset, with and without batch integration. Figure 2 shows a graphical abstract illustrating the overall workflow of the methods used for single cell multi-omics integration in this work. Additionally, UMINT has been validated on a bulk multi-omics dataset for integration and classification.

**Figure 2:**
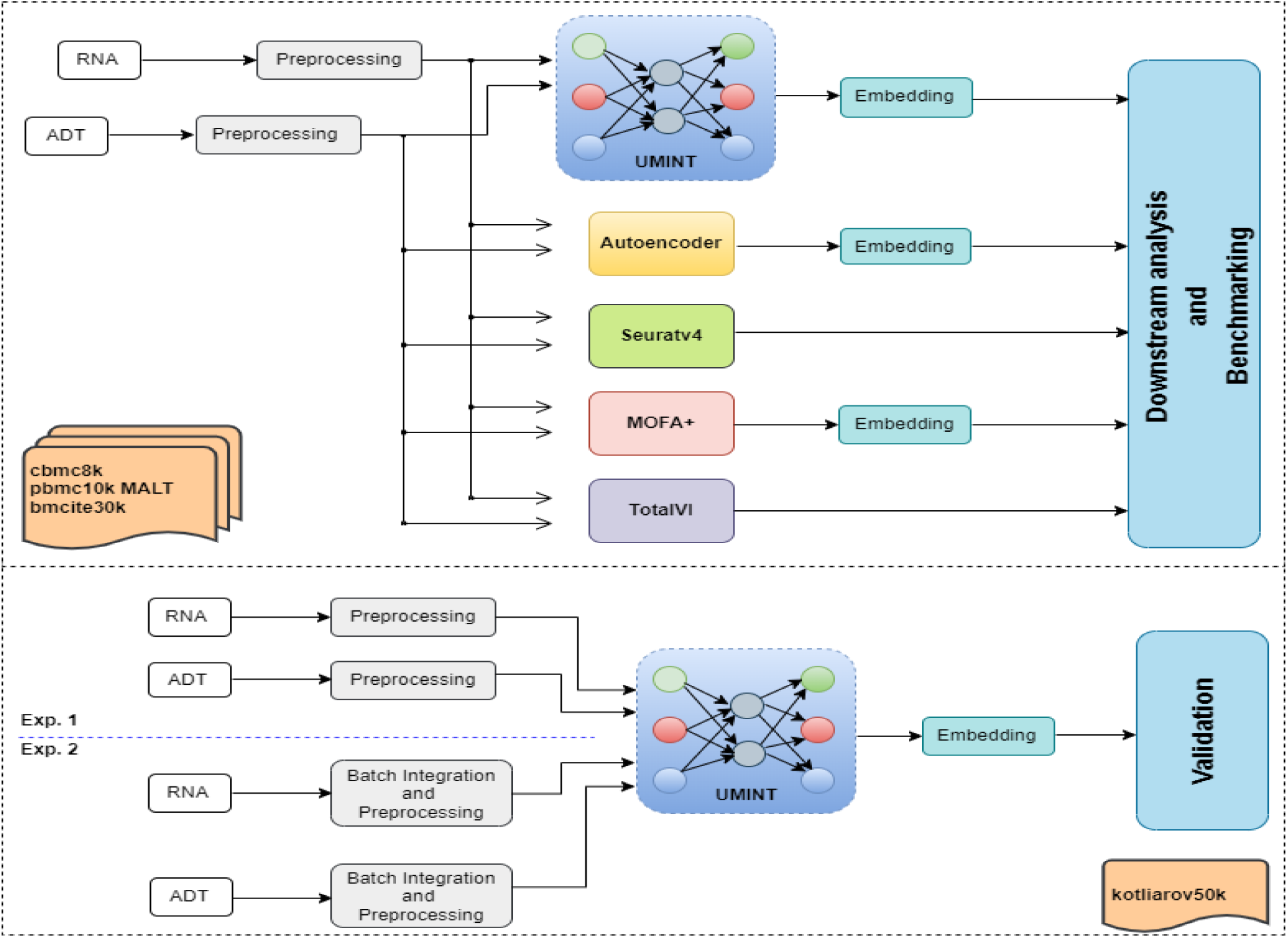
A graphical abstract showing the overall workflow of the methods used for evaluation and benchmarking of UMINT single cell multi-omics integration.

### 3.1 Comparison with other unsupervised neural network models

The architecture of UMINT is similar to that of an autoencoder (AE) network, which is also an unsupervised neural network used for dimension reduction [30]. AE being the closest resemblance to UMINT, we have first compared it with AE, both in terms of architectural difference and performance.

#### 3.1.1 Comparison with AE based on number of trainable parameters

Similar to an autoencoder network, UMINT also tries to reconstruct the original input as discussed in Section 2.2. However, there is a few differences between the two. The input layer in AE shares a dense connection with the first hidden layer, whereas, the connections between the *Input layer* and the first *Modality encoding layer* in UMINT is not dense. This reduces the number of parameters to be trained drastically. Although, in this work, UMINT has been used to integrate single cell RNA and protein expression data, it is quite capable of integrating any number of omics layers. Let us consider that the input to UMINT consists of data from *m* modalities having n samples each with *d*_1_, *d*_2_, …, *d_m_* dimensions respectively. As shown in Figure 1, UMINT consists of the same number of modules in the *Modality encoding layer* as that in the *Input layer.* If the number of neurons in each of the module of this *Modality encoding layer* are *n*_1_, *n*_2_,…,*n_m_* respectively, then the total number of trainable parameters (*TP_UMINT_*) between the *Input layer* and the *Modality encoding layer* in the encoder network becomes

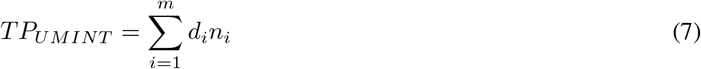

Considering an input of similar dimensions, if an AE network is employed to achieve this same task of integration, the number of trainable parameters (*TP_AE_*) between the input layer and the first hidden layer becomes

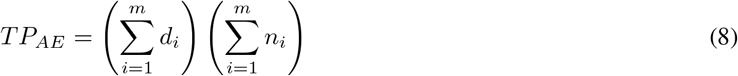

Thus, the reduction in the number of trainable parameters (*TP_Reduction_*) in the encoder network is given by

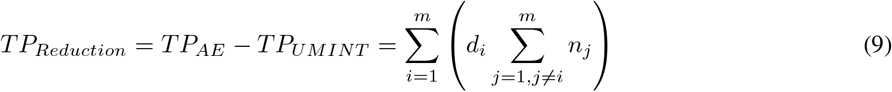

A reduction in the number of trainable parameters by the same amount is also available at the *Reconstruction layer* of the decoder network. Hence, the total reduction (*TP_TotalReduction_*) in the number of trainable parameters in UMINT is given by

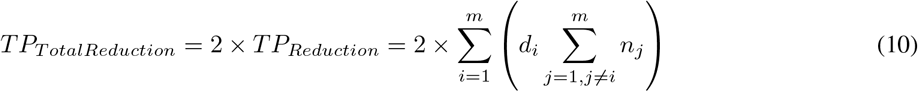

This is a massive improvement over a regular AE network and makes UMINT computationally far less expensive.

#### 3.1.2 Comparison with AE based on performance

Initially, the performance of UMINT has been compared with that of an AE with respect to their reconstruction capability and the latent embedding of reduced dimensions produced at the bottleneck layer of each network. The RNA and protein expression data have been stacked together to form the input to the AE. Keeping the hyper-parameter values identical, we have made the autoencoder reconstruct the input by passing it through a series of layers. In the process, it learns to extract useful features at the bottleneck layer. In a similar fashion, we have reconstructed the RNA and the protein expression data using UMINT, and computed Pearson correlation coefficient between the original data and its reconstructed counterpart. As shown in Figure 3, we have observed that UMINT has been able to reconstruct RNA and ADT efficiently for all the datasets. The reconstruction of RNA is closely comparable with that achieved using AE, while in case of ADT, UMINT has produced far more accurate reconstruction than AE. Stability of the results have been ensured by repeating the experiments 10 times.

**Figure 3:**
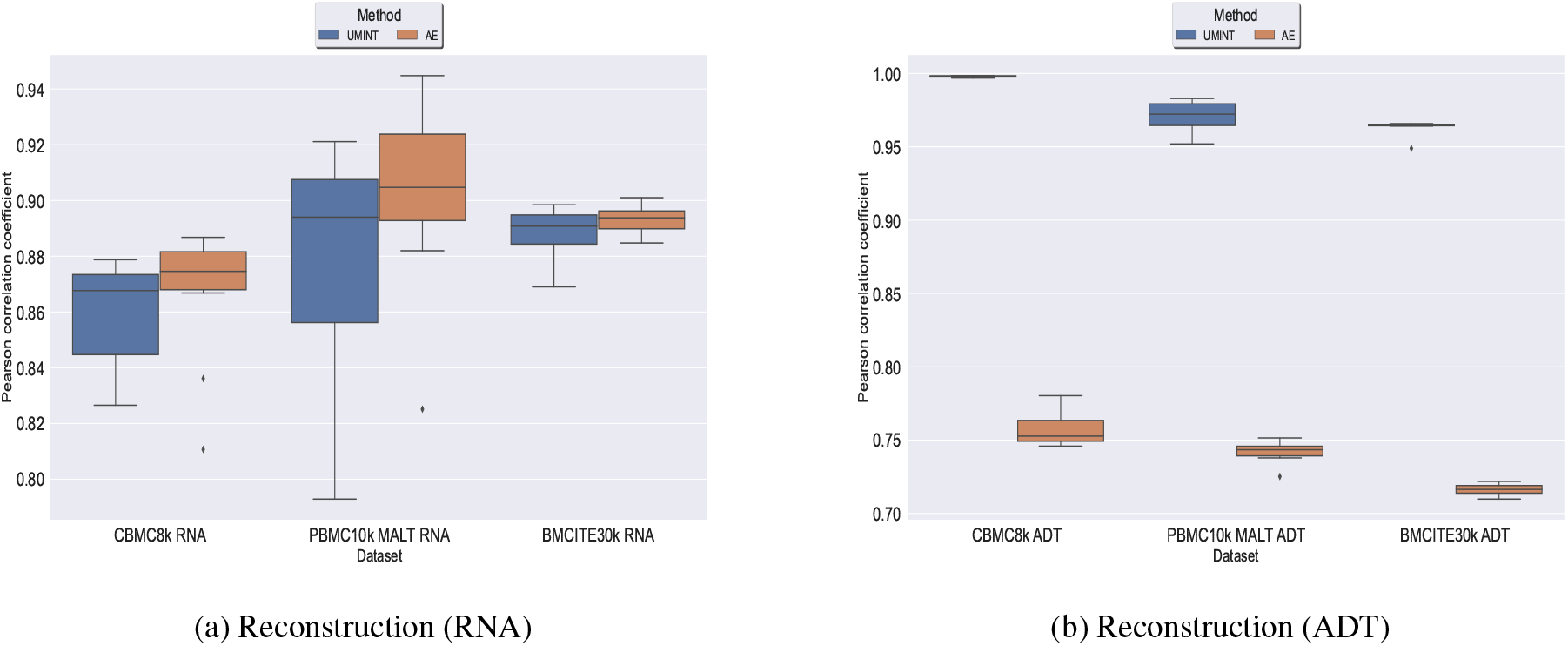
Comparison of reconstruction performance of UMINT with that of an AE

Thereafter, we have compared the latent low-dimensional embedding produced by UMINT with that produced by an AE. Once trained to create a lossy reconstruction of the input, the latent representation has been extracted from the bottleneck layer of both UMINT and AE. Cell-type clustering on this latent embedding using k-means and agglomerative hierarchical clustering has been performed to validate the effectiveness of UMINT and compare it with AE. Adjusted Rand Index (ARI) and Fowlkes Mallows Index (FMI) scores have been used to measure the degree of agreement between the actual and predicted cell-types.

For two sets of cluster labels, the overlap between them is represented by a contingency table **C** = |*c_ij_*], where *c_ij_* indicates the total number of points belonging to both *i^th^* cluster of the first set and *j^th^* cluster of the second set. ARI is an external cluster validity index, and is thus defined as

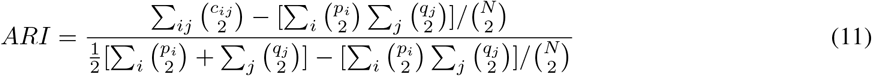

where *p_i_* = ∑_*j*_ *c_ij_*, *q_j_* ∑_*i*_ *c_ij_* and *N* = ∑_*ij*_ *c_ij_* respectively. An ARI value close to 1 indicates good resemblance between two clusters. Similarly, FMI, another external evaluation index used to measure the similarity between two sets of cluster labels, is defined as

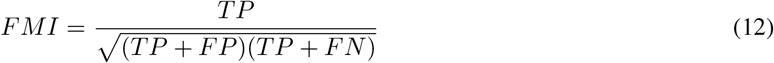

where *TP*, *FP* and *FN* denotes the count of True Positives, False Positives and False Negatives respectively. The FMI score lies between 0 and 1, and a high value implies a good similarity between two clusters.

We have observed that for *cbmc8k* and *bmcite*30*k* datasets, the latent representation produced by UMINT has been more representative of the cell clusters when compared with that of AE. This has been validated by both ARI and FMI scores as shown in Figure 4. For *pbmc*10*kMALT*, when hierarchical clustering algorithm is used, UMINT embedding has produced similar ARI scores to that obtained on AE embedding (with comparable median ARI scores) but poor FMI scores than that obtained on AE embedding. On applying k-means clustering algorithm, UMINT embedding has, however, produced better ARI scores than AE embedding and similar FMI scores as compared to that obtained on AE embedding (Figure 4). All experiments have been repeated multiple times to validate stability of the results.

**Figure 4:**
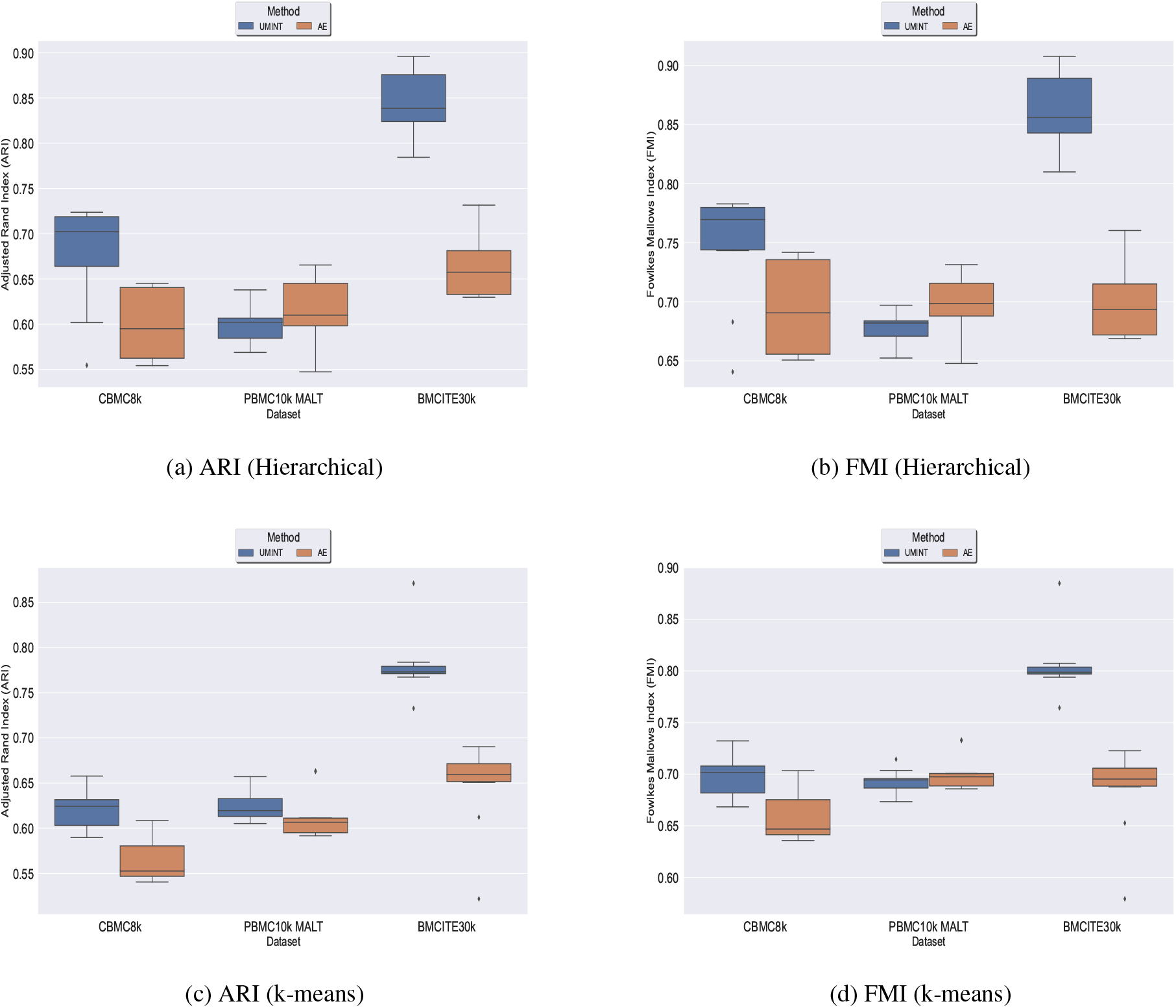
Comparison of clustering performance of UMINT against that of AE when agglomerative hierarchical clustering is used, as measured by (a) ARI and (b) FMI; (c) and (d) show clustering performance of UMINT compared to AE, as measured by ARI and FMI respectively, when k-means clustering algorithm is used.

### 3.2 Comparison with other state-of-the-art methods for single cell multi-omics integration

Subsequently, we have compared the performance of UMINT with three other state-of-the-art methods - Seuratv4 [16], MOFA+ [17] and TotalVI [18]. We have chosen these three methods since they represent three different categories of algorithms - Graph based, Matrix factorization based and Neural network based, developed for single cell multi-omics integration [24]. To ensure a fair comparison between all these methods, we have followed the same preprocessing pipeline for all the datasets used for comparison. The effectiveness of UMINT has once again been demonstrated by clustering the cell-types on the latent low-dimensional embedding produced by it. It may be mentioned here that Seuratv4 is capable of producing an integrated low-dimensional representation through weighted-nearest neighbor analysis, and also find cell clusters from the integrated embedding using Louvain [31], Leiden [32] or SLM [33] community-detection algorithms. TotalVI, on the other hand, integrates the data through variational inferencing and autoencoding, and uses the standard Scanpy [34] pipeline for clustering on the latent embedding. MOFA+, however, only produces a low-dimensional representation of the integrated data, as in the case of UMINT. The methods used in this work for comparison have been compared theoretically in Table 2. Thus, for Seuratv4 and TotalVI, we have extracted the cluster labels, while for MOFA+, we have extracted the factors representing the low-dimensional embedding and used hierarchical and k-means clustering on the same, similar to UMINT, as illustrated in the graphical abstract shown in Figure 2. Interestingly, UMINT has outperformed all three methods in terms of ARI and FMI scores, validated both by hierarchical and k-means clustering. Figure 5 shows the average ARI and FMI scores obtained using UMINT plotted against the scores obtained by the three benchmark methods.

**Figure 5:**
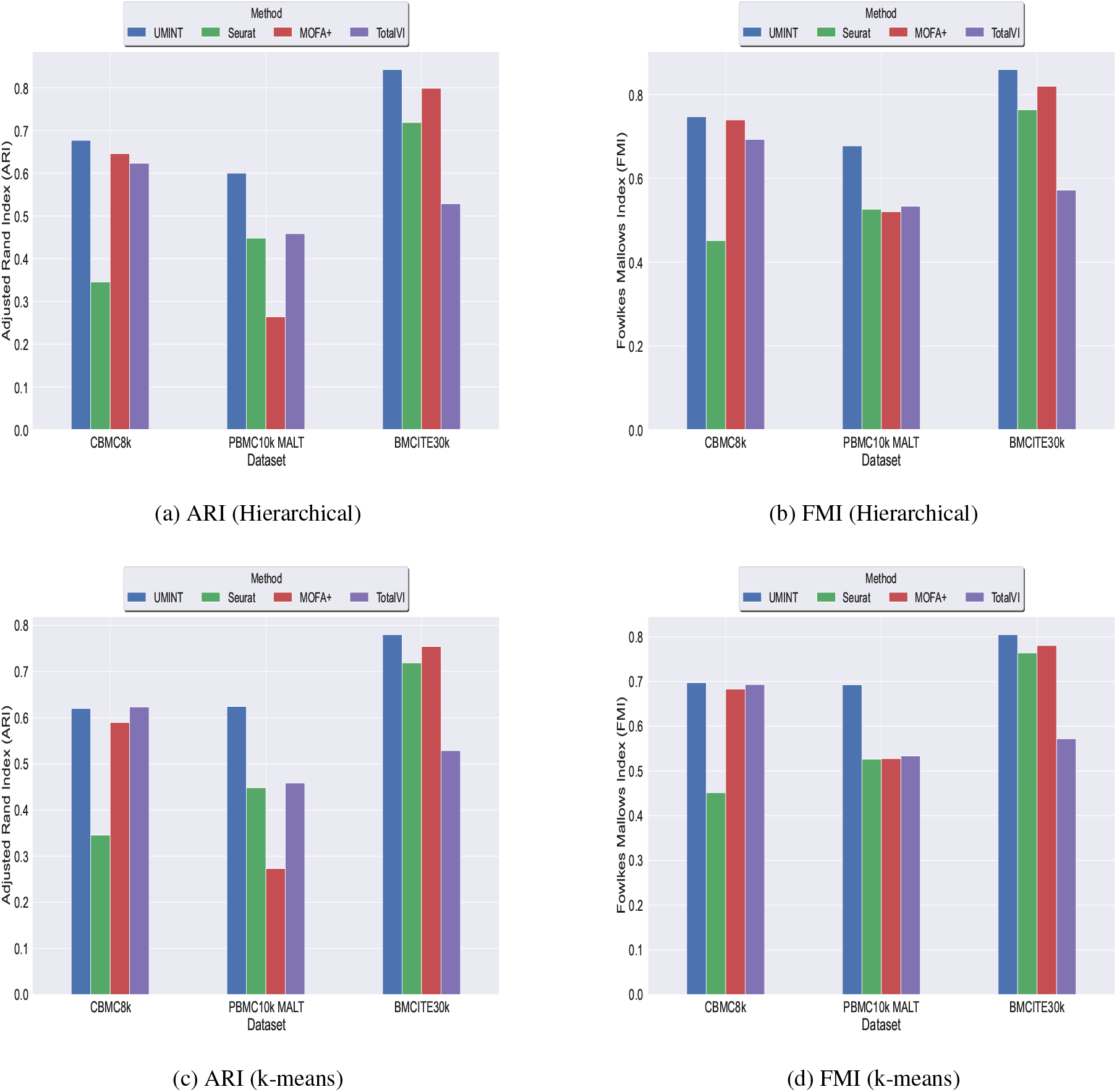
Comparison of clustering performance of UMINT against Seuratv4, MOFA+ and TotalVI when agglomerative hierarchical clustering is used, as measured by (a) ARI and (b) FMI; (c) and (d) show performance of each method, as measured by ARI and FMI respectively, when k-means algorithm is used for clustering.

**Table 2:** A theoretical comparison between UMINT and other methods used for comparison

| Method | Methodology | Produces latent embedding (Yes/No) | Support cell-type clustering (Yes/No) | Omics integration supported for | Makes assumption about data distribution (Yes/No) | Can reconstruct original data (Yes/No) |
| --- | --- | --- | --- | --- | --- | --- |
| UMINT | Neural network based | Yes | No | Both single cell and bulk | No | Yes |
| Autoencoder | Neural network based | Yes | No | Both single cell and bulk | No | Yes |
| Seuratv4 | Graph based | Yes | Yes, via Louvain, Leiden and SLM | Single cell only | No | No |
| MOFA+ | Matrix factorization based | Yes | No | Both single cell and bulk | Yes | Yes |
| TotalVI | Neural network based | Yes | No (recommends using Scanpy) | Single cell only | Yes | Yes |

### 3.3 Performance of UMINT on multi-batch datasets

Batch effects in single cell datasets pose great challenges in data integration and compromises the results [35, 36]. We wondered how UMINT would perform when there are batches in the data. The dataset *bmcite*30*k* used in this work contains two batches. However, we have not performed batch integration on this dataset. Figures 3 - 5 show the performance of UMINT on *bmcite*30*k* dataset when no batch integration has been performed. We have further observed that batches present in the *bmcite30k* projection by UMINT have been well integrated and are thus inseparable. Additionally, cell clusters obtained on UMINT projection are cohesive and well separated too. Thus, we can conclude that besides cell-type clustering, UMINT has been able to correct batches present in *bmcite*30*k* dataset efficiently. The results in support of this claim have been shown in Figure 6.

**Figure 6:**
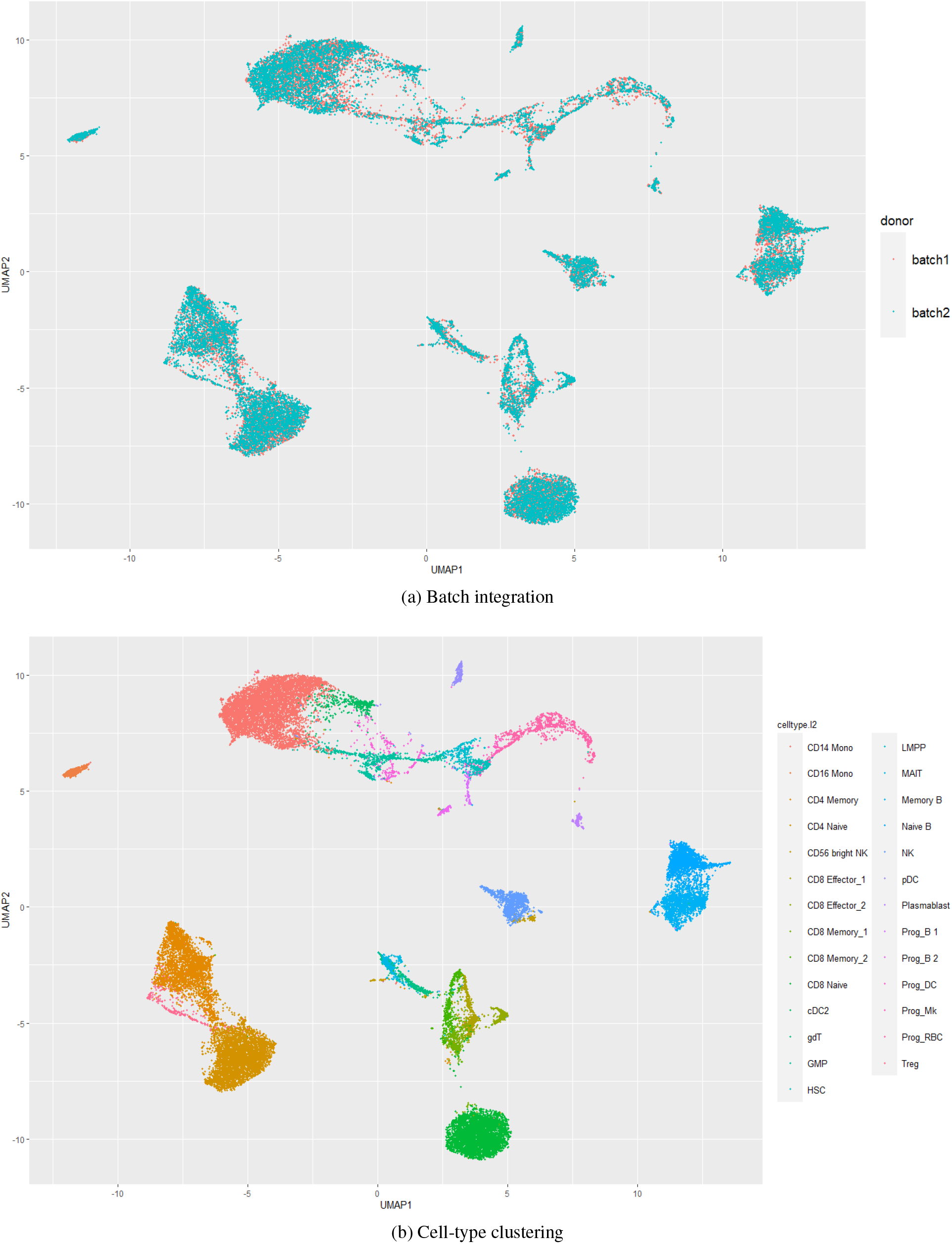
Performance of UMINT on *bmcite*30*k* dataset with respect to (a) batch integration, (b) cell-type clustering

In order to reinforce our findings, we have performed a few more experiments on another dataset *kotliarov*50*k.* This dataset, collected from [19], contains filtered data for 52117 cells with highly variable genes (3999), and two batches of RNA and protein expressions each. Moreover, it contains expression values for 87 proteins, a lot more than the three other datasets used in this work. We have first integrated batches using Seuratv4 SCTransform() [16] module with default parameters and fed the batch integrated RNA and ADT datasets to UMINT. The low-dimensional embedding produced by UMINT has then been evaluated for clustering and batch integration performance. In another experiment, we have fed the preprocessed RNA and ADT datasets (without batch integration) into UMINT, and evaluated the low-dimensional embedding produced by it for clustering and batch integration performance too. Apart from two external validity indices, we have used two internal validity indices - silhouette coefficient [37] and Davies Bouldin (DB) index [38] to measure the clustering performance of the UMINT-generated embeddings on the omics data with and without batch integration. Interestingly, we have observed that when the UMINT embedding has been generated from the RNA and ADT data without batch integration, the ARI, FMI, Silhouette and DB scores achieved have been quite close to those achieved when UMINT embedding has been generated from batch integrated RNA and ADT data, as shown in Figure 7. Thus, it is clear that even without batch integration, UMINT can extract most relevant features from the data that can act as input to further downstream investigations. However, the UMINT-generated embedding obtained on batch integrated data has shown better batch correction performance than that obtained on data without batch integration. From Figure S1 (in Supplementary Material), it can be observed that batches in *kotliarov*50*k* data remain separable if batch integration is not performed on the dataset explicitly. This explains why performance on *kotliarov*50*k* dataset without explicit batch correction is not as good as performance on the same dataset with batch correction. Thus, there is further scope of improvement for UMINT in terms of batch correction performance.

**Figure 7:**
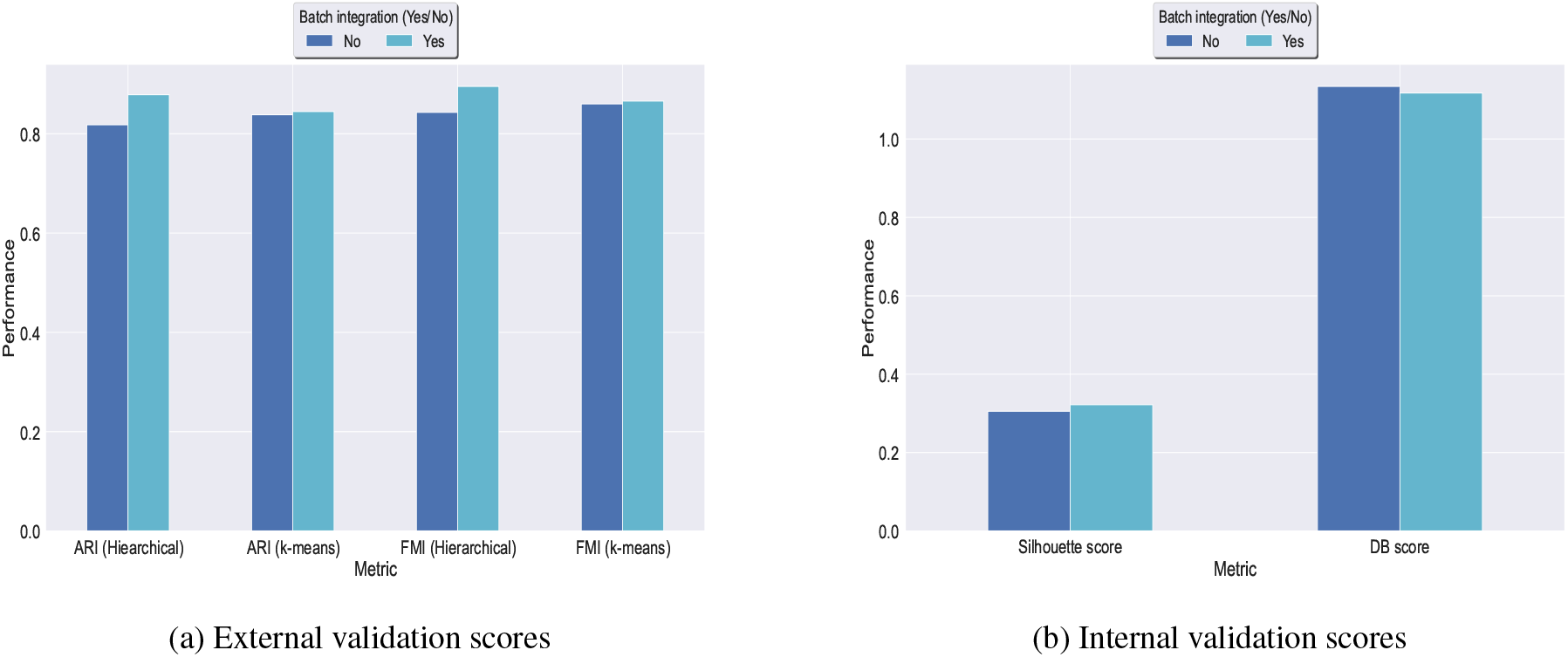
Clustering performance of UMINT-generated embeddings obtained from RNA and ADT data with and without batch integration on the *kotliarov*50*k* dataset as measured by (a) external validity indices and (b) internal validity indices.

### 3.4 Performance of UMINT on bulk multi-omics

Finally, UMINT has been assessed for its integration performance on bulk expression datasets. The TCGA multi-omics data for Liver Hepatocellular Carcinoma (LIHC) from the TCGA portal (now moved to Genomic Data Commons https://gdc.cancer.gov/), has been used for this purpose. The dataset contained three omics layers - DNA methylation (DNAm), Copy Number Variation (CNV) and RNA-seq. Pre-processed datasets collected from [39], contains 404 paired samples out of which 359 are cancer and 45 are normal.

To assess the effectiveness of UMINT on bulk multi-omics data, we have first compared the reconstruction and classification performance of UMINT against that obtained using an AE. As shown in Figure S2 (in Supplementary Material), UMINT has been able to reconstruct DNAm and RNA-seq better than AE, while reconstruction for CNV compares favourably with that obtained using AE. For assessing the classification performance, we have trained UMINT on 80% data and used the remaining 20% as test data. The overall classification accuracy, recall and F1-scores achieved by UMINT projection have been higher than that achieved by AE (Figure S2 in Supplementary Material).

In our earlier work [39], we had proposed a deep denoising autoencoder and multi-layer perceptron (DDAE-MLP)-based method for integrating such bulk multi-omics datasets, though the question addressed earlier was whether we could predict one omics data (gene expression) from the other two (DNAm and CNV), in case the mRNA data is of degraded quality or not available at all. As already stated in Section 3.1.1, UMINT can efficiently integrate variable number of omics layers. In order to explore this further, in this work, we have tried to find out if UMINT can capture better feature variability by integrating all three omics layers (DNAm, CNV and RNA-seq) simultaneously. Additionally, since MOFA+ can integrate both single cell and bulk multi-omics data, we have considered MOFA+ while comparing the classification performance of UMINT as well. With the same 80 - 20 train-test split, UMINT has been observed to perform competitively against MOFA+, as measured by classification accuracy, precision, recall and F1-scores. Figure S3 (in Supplementary Material) shows the results for this part of evaluation.

## 4 Discussion and Conclusion

In this work, we have introduced a novel deep Unsupervised neural network for single cell Multi-omics INTegration (UMINT). We have used UMINT to integrate heterogenous single cell omics modalities and extract meaningful projections at reduced dimensions. These features have been further used for clustering. The effectiveness of UMINT has been evaluated on four publicly available CITE-seq datasets, and compared on three of them with some other state-of-the-art algorithms used for multi-omics integration.

The strengths or advantages of UMINT are many-fold. UMINT-generated latent embedding has been proved to produce better clustering as compared to other existing unsupervised neural network model, like AE. UMINT-generated reconstructions have been better than or at least as good as that produced by AE, and that too using far less number of parameters. When evaluated against other benchmark algorithms, UMINT has displayed superior performance over every other method used for comparison, across most of the evaluation criteria on all the three datasets. Moreover, UMINT does not make assumptions about the underlying data distribution, thus making it more robust. Finally, UMINT has been found to be competent enough to integrate bulk multi-omics datasets too. It has been able to produce better or comparable reconstructions for bulk omics data like that obtained using AE. UMINT-extracted features have also been effective enough in classifying tumour and normal samples, which also implies that UMINT supports integration of variable number of omics modalities. Very few such integration methods exist that can efficiently integrate features from both single-cell and bulk multi-omics, and can handle variable number of omics layers.

UMINT, however, is susceptible to batch effects to some extent. It has been able to correct batches for *bmcite*30*k* dataset perfectly, while for *kotliarov*50*k* data, integration has been compromised by a tolerable amount due to batch effect. Addressing this shortcoming remains as a future extension to this work. In the current scope of work, we have not explored the integration of other single cell omics modalities. Such integration would further allow us to better understand the overall contribution of epigenomics layer at a single cell level in regulatory systems biology on top of scRNA-seq and protein expression data. This also remains a future extension to UMINT.

Nevertheless, UMINT can capture better variability among high-dimensional datasets and produce robust lowdimensional embedding which would assist in further downstream analyses. A reduction in the number of trainable parameters also makes UMINT far less computationally expensive that other existing neural network models. Thus, we are able to provide a robust and efficient unsupervised deep learning model for single-cell multi-omics integration.

## Data and Code availability

UMINT has been implemented in Python 3. The codes to reproduce the results are freely available at https://github.com/deeplearner87/UMINT. The preprocessed datasets used in this work can be downloaded from https://doi.org/10.5281/zenodo.6349408.

## Author’s contribution

Conceptualization of methodology and framework: CM, DBS, VD, RKD. Data Curation, Data analysis, Formal analysis, Visualization, Implementation: CM, DBS. Investigation, Code review, Validation, Original draft preparation: DBS, CM, VD. Reviewing, Editing, Overall Supervision: RKD.

## Acknowledgements

DBS thanks the Data Science Laboratory, A. K. Choudhury School of Information Technology, University of Calcutta, Kolkata, India for providing the HPC infrastructure to execute and test some of the models. RKD acknowledges SyMeC Project grant [BT/Med-II/NIBMG/SyMeC/2014/Vol. II] given to the Indian Statistical Institute by the Department of Biotechnology (DBT), Government of India.

