## Supplementary material for "UMINT: Unsupervised Neural Network For Single Cell Multi-Omics Integration"

Chayan Maitra[1,*], Dibyendu Bikash Seal[2,*], Vivek Das[3], and Rajat K. De[4,**]

[1,4]*Machine Intelligence Unit, Indian Statistical Institute, 203 Barrackpore Trunk Road, Kolkata 700108, India.*
[2]*A. K. Choudhury School of Information Technology, University of Calcutta, JD-2, Sector - III, Saltlake, Kolkata - 700106, India.*
[3]*Novo Nordisk A/S, Novo Nordisk Park 1, 2760 Maløv, Denmark.*
[*]*Authors contributed equally*

April 21, 2022

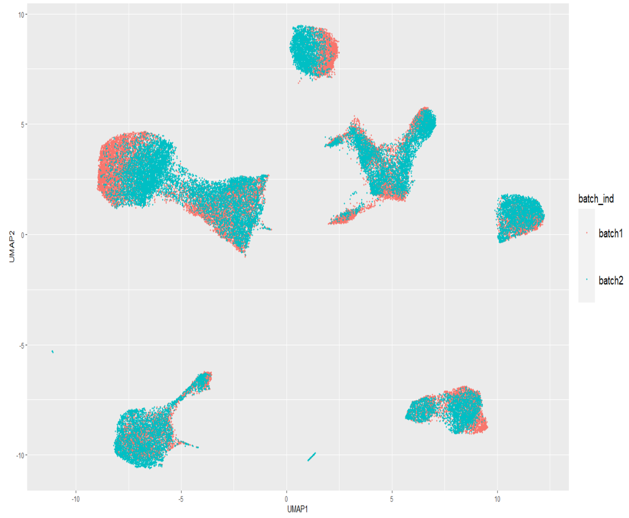


(a) Batch correction performance of UMINT on $kotliarov50k$ dataset without batch integration

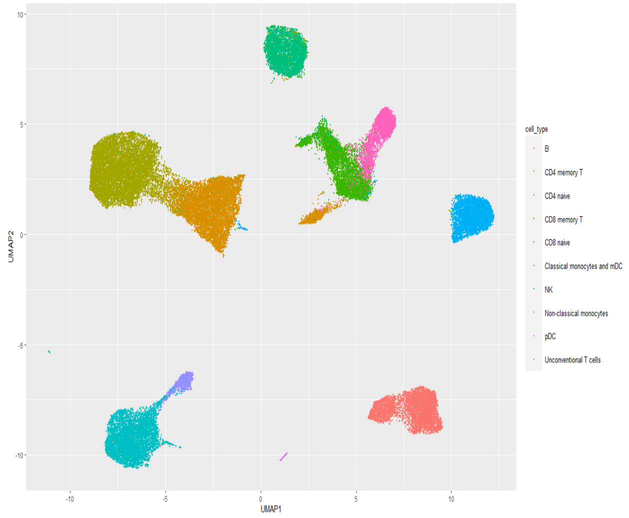


(b) Cell-type clustering performance of UMINT on $kotliarov50k$ dataset without batch integration

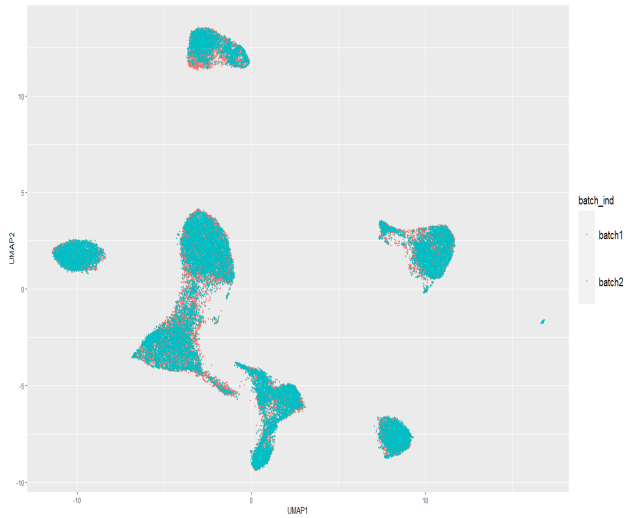


(c) Batch correction performance of UMINT on $kotliarov50k$ dataset with batch integration

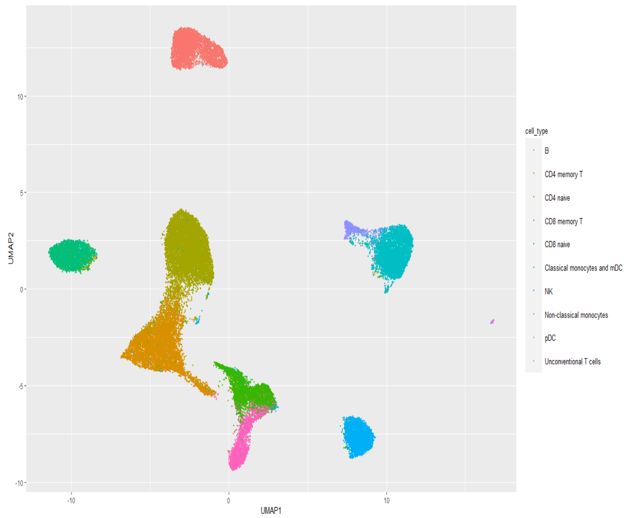


(d) Cell-type clustering performance of UMINT on $kotliarov50k$ dataset with batch integration

Figure S1: (a) and (b) show batch correction and clustering performance of UMINT on the $kotliarov50k$ dataset without batch integration; (c) and (d) show similar results on the $kotliarov50k$ dataset with batch integration

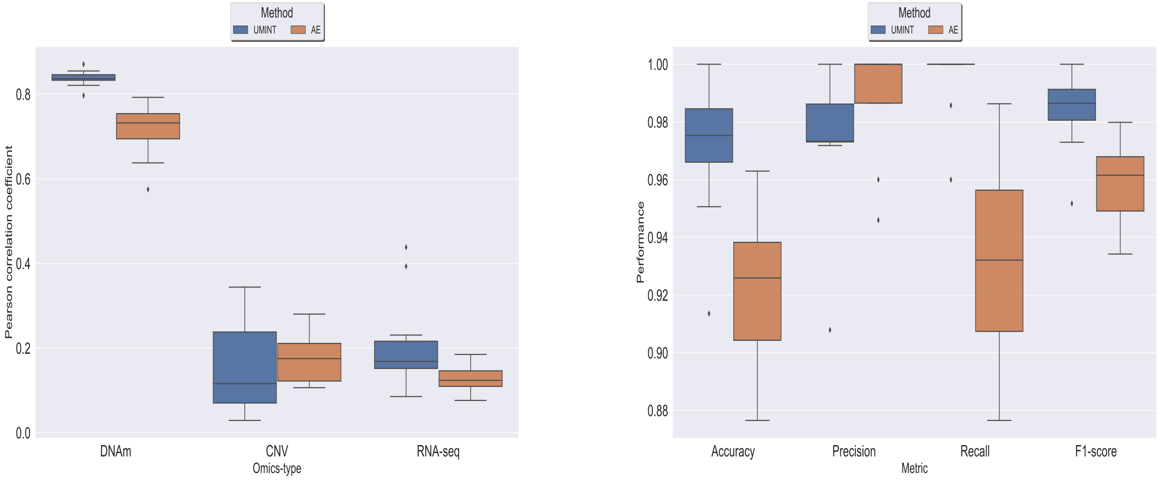


(a) Reconstruction (b) Classification of Tumour/Normal Samples

Figure S2: (a) shows the performance of UMINT compared against an autoencoder (AE) network with respect to reconstruction of DNA methylation (DNAm), Copy Number Variation (CNV) and RNA-seq expressions for TCGA LIHC dataset; (b) shows the performance of UMINT compared against an AE with respect to classification of tumour and normal samples on the TCGA LIHC dataset

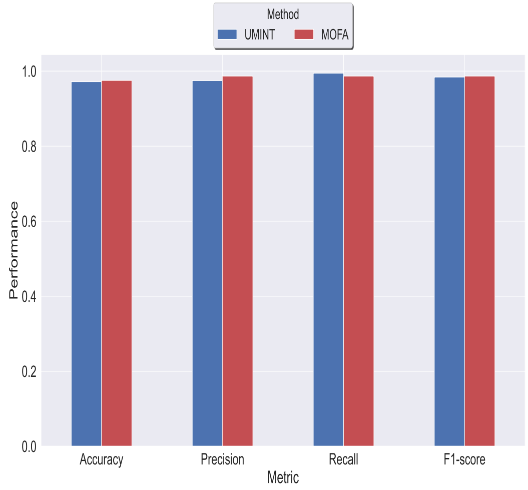


Figure S3: Classification performance achieved on UMINT-generated low-dimensional embedding compared against that obtained on MOFA+-generated embedding for the TCGA LIHC dataset.
